# Spatial trophic cascades in communities connected by dispersal and foraging

**DOI:** 10.1101/469486

**Authors:** David García-Callejas, Roberto Molowny-Horas, Miguel B. Araújo, Dominique Gravel

**Affiliations:** CREAF, Cerdanyola del Vallès, 08193, Spain; Departamento de Biogeografía y Cambio Global, Museo Nacional de Ciencias Naturales, Consejo Superior de Investigaciones Científicas (CSIC), Calle José Gutiérrez Abascal 2, 28006 Madrid, Spain

## Abstract

Pairwise interactions between species have indirect consequences that reverberate throughout the whole ecosystem. In particular, interaction effects may propagate in a spatial dimension, to localities connected by organismal movement. Here we study the propagation of interactions with a spatially explicit metacommunity model, where local sites are connected by dispersal, foraging, or by both types of movement. We show that direct and net effects of pairwise interactions may differ in sign when foraging across localities is prevalent. Further, the effect of a species over another in the local community does not necessarily correspond to its effect at the metacommunity scale; this correspondence is again mediated by the type of movement across localities. Networks of net effects are fully connected, indicating that every species in the metacommunity has a non-zero influence on every other species. Lastly, the magnitude of net effects between any two species strongly decays with the distance between them. These theoretical results strengthen the importance of considering indirect effects across species at both the local and regional scale, point to the differences between types of movement across locations, and thus open novel avenues for the study of interaction effects in spatially explicit settings.

## Introduction

Ecological communities are complex systems in which species interact with each other through a multitude of pathways. The effect of a species on the rest of the ecosystem is generally difficult to predict and quantify, and likely depends on factors such as the number and magnitude of interactions in which it engages (Zhao et al., 2016), or the structure of the overall network. Network topology that may ultimately enhance or buffer the propagation of the initial direct effects (Polis, 1994). Trophic cascades are striking examples of interaction effects propagating through food chains: changes in the occurrence, strength or outcome of a certain trophic interaction often have a significant top-down influence on the rest of the community (Schmitz et al., 2000).

Just as the spreading of disease (Balcan et al., 2009) or information (Barthélemy, 2011) in other types of complex networks, the propagation of interaction effects across ecological networks has an obvious spatial dimension, as interaction cascades often link organisms that are spatially disconnected. Thus, we may define a *spatial cascade* as a set of indirect interactions that spread in a spatial dimension, potentially linking disconnected species. Spatial cascades are ubiquitous in nature: they connect for instance different regions by migratory animals (Springer et al., 2018) or, on a more local scale, separated locations by dispersing individuals (Leibold et al., 2004). Spatial cascades may occur between different locations of a single habitat type. For example, predator species may consume bird eggs from nests of different forest patches (Chalfoun et al., 2002), with potential feedbacks for the bird populations and associated resources. The connections across different habitats by flows of nutrients or organisms have also been well documented (Polis et al., 1997). Focusing on the flow of organisms, just to note two prominent examples, Estes et al. (1998) documented how otter predation by killer whales in the open North Pacific triggered an increase in the biomass of sea urchins in the nearshore habitat of the Aleutian archipelago, ultimately driving a strong decline in kelp density. More recently, Knight et al. (2005) showed that the presence of predatory fish in ponds reduced the number of adult dragonflies in the surrounding area, which resulted in a significant increase in pollinator density and subsequent reproductive success of terrestrial plants, as compared to areas close to ponds without fish predation.

Both bottom-up and top-down effects may be involved in spatial cascades, and these effects will likely differ in magnitude depending on the type of spatial flux or the trophic level of the connecting species (Allen and Wesner, 2016). For example, variations in the magnitude of plant dispersal between neighbouring locations will trigger bottom-up community-level responses on all sites (Christian, 2001). On the other hand, predators foraging on spatially disconnected patches may induce top-down indirect effects that may propagate across patches, either through consumptive effects that spread down the local trophic chains (Polis et al., 1997) or by non-consumptive effects on prey species (Orrock et al., 2008). Overall, despite the growing number of studies documenting spatial propagation of interaction effects, the concept of spatial cascades has not yet been rigorously explored and generalized. For example, there are currently no theoretical hypothesis on the decay of the magnitude of net effects with spatial distance, or on whether different modes of movement generate similar or different patterns of effect propagation.

The net interaction effect between any two interacting species is, conceptually, the sum of their direct effects from pairwise interactions and indirect effects mediated by other species or entities (Abrams, 1987). The direct effects of a species over another can be formulated in several ways (Berlow et al., 2004), but generally involve the effects over some property of interest at the population level, such as short-term growth rate (Abrams, 1987). Indirect effects, in turn, involve all effects between two species that do not occur via direct interactions. Indirect effects may occur between species that interact directly or not, via the propagation of effects over the ecological network. These effects have been classified as being triggered by changes in the abundance of the intermediary species (*density-mediated indirect interactions*) or by these intermediary species modifying the context of a direct interaction (see e.g. Wootton 2002 for further definitions and examples). It has been repeatedly shown that indirect effects may be as strong, or even stronger than direct effects, up to the point of switching interaction net effects from positive to negative or viceversa (e.g. Menge 1995).

The metacommunity concept (Leibold and Chase, 2018) provides a comprehensive theoretical framework for studying the propagation of interaction net effects in a spatially explicit setting. In virtually all metacommunity studies we are aware of, it is assumed that species connect the local communities via dispersal, i.e. the permanent establishment of individuals on a different territory from their birthplace. Dispersal, however, is not the only process by which species can link spatially disconnected patches. Foraging, the active search for food of a mobile individual, may link the trophic chain of its reproductive area with other, potentially disconnected communities in which the individual acquires varying fractions of its diet (McCann et al., 2005). If foraging species are based on a central site, which may well be their reproductive area, their foraging effort and associated effects on local communities generally decay with distance, in what is termed *Central-place foraging*, (Orians, 1979). Just as with dispersal, the spatiotemporal dynamics of foraging are extremely varied, with variation in home ranges spanning several orders of magnitude (as an indicator, Swihart et al. 1988 collected data on 23 species of mammals whose home range varied from 0.05 to 2285 ha). Of course, both foraging and dispersal modes of movement occur in nature and are not independent from each other, but as a first approximation, we may expect spatial cascades triggered by each movement type to display different properties and effects on the connected local communities (Fig. 1). For example, in two simple food chains connected by a dispersing species, the net flow of individuals from one community to the other will benefit the predators of the dispersing species, and in turn, adversely affect its prey. On the other hand, if the same species connects the two food webs by foraging sporadically on the second location, it will trigger a negative effect up the trophic chain of that location, and will benefit species on which the preyed species feeds. Here we study how net effects are propagated in space when local food webs are connected by dispersal, foraging, or a combination of both movement types, using model multi-trophic metacommunities. In particular, we ask the following questions: (1) What is the distribution of signs and magnitudes of net effects in communities connected by dispersal, foraging, or a mixture of both? (2) Do metacommunities connected by different movement modes vary in their topological structure? (3) Does the magnitude of the net effects between any two species decay with increasing distance between them?

**Figure 1:**
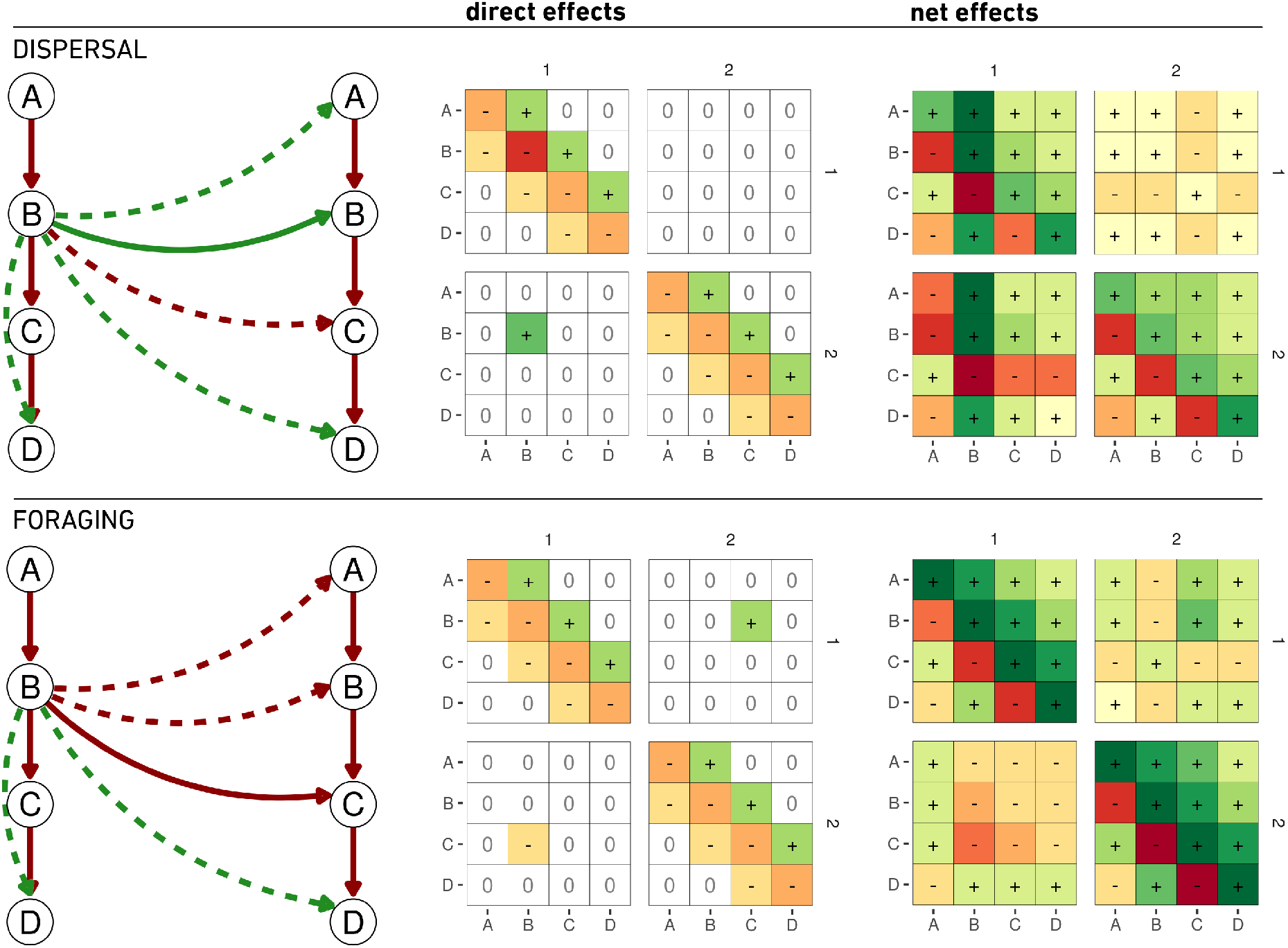
Interaction matrices and net effect matrices for two simple configurations. In both cases, a linear food chain is represented at two locations (*1* and *2*). In the dispersal configuration, species B disperses from the first location to the second. In the second configuration, species *B* preys on species *C* on both locations. Green links represent positive effects, red negative. Solid arrows represent direct effects, dashed arrows expected indirect effects. For clarity, in the food chains we display only the indirect effects arising directly from species *B* at location *1*. Darker shades in the matrices indicate stronger effects. The matrices can be read as with the following example: in the foraging configuration, the direct effect of *B* in location *1* over *C* in location *2* is given by locating the column that indicates species *B* at location *1* (the second column of the matrix), and the row indicating species *C* at location *2* (seventh row).

## Methods

We developed a spatially explicit metacommunity model in which local trophic communities are connected through 1) dispersal, 2) foraging, or 3) both. The dynamics of the system are given by a general Lotka-Volterra implementation, following Gravel et al. (2016), and for each configuration we ran numerical simulations and recorded both the direct effect and the net effect between each pair of species in the metacommunity, as well as a set of network metrics for characterizing potential differences in metacommunity structure.

### Quantification of direct and net effects

In theoretical analyses of ecological networks, the Jacobian matrix of the system (also called *community matrix*) is widely used to describe the direct effects between each pair of species at equilibrium. In its most common implementation, it represents the effect on one species’ growth rate in response to small changes in another species’ abundance (Berlow et al., 2004; Novak et al., 2016). Consider a general population dynamics model of *S* species in which the population density of species *i* over time is given by

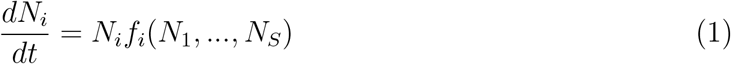

where *N_i_* is the density of species *i*, and *f_i_*(*N*_1_,…, *N_S_*) is its growth rate, potentially influenced by any other species. In this general case, the elements of the Jacobian matrix *C* are

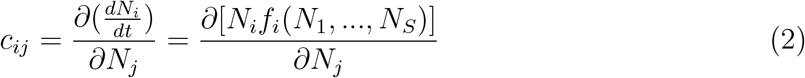

The net effect of species *j* over species *i*, in turn, is the sum of its direct effects and all indirect effects between the two species (Bender et al., 1984; Montoya et al., 2009; Novak et al., 2016). The net effect matrix of a community is defined as the negative of the inverse Jacobian matrix, i.e. −*C*^−1^, and its coefficients represent the net effect of an increase in species *j*’s population growth rate on the density of species *i*, when all species respond to direct effects (Novak et al., 2016).

### The model

The dynamics of the community are modelled with a general Lotka-Volterra implementation, following Gravel et al. (2016). Considering a set of *S* species present at *n* locations, the dynamics of species *i* at location *x* is given by:

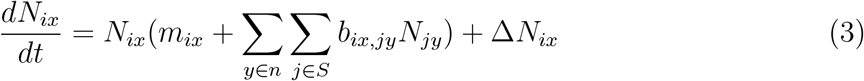

where *m_ix_* is the intrinsic growth rate of species *i* at location *x*, *N_ix_* its abundance, Δ*N_ix_* is the net migration balance, and *b_ix,jy_* is the per capita effect of species *j* at location *y* on species *i* at location *x*. The parameter *b_ix,jy_* encapsulates the effect of foraging to/from other locations, and it represents a basic situation in which species *i* moves out of its reproductive area *x* to feed at location *y* on species *j*. Specifically, we assume that a foraging species allocates a fraction *f* of its foraging efforts to communities outside its reproductive location, which implies that the effort allocated to feeding in its local community is 1 *− f*. We incorporate this in our model as follows:

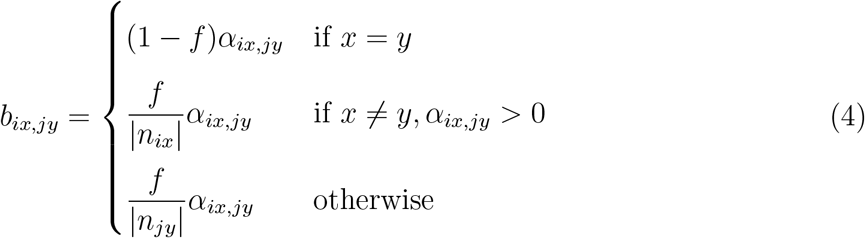

The first situation corresponds to the effect of predation from the same location, in which case the interspecific interaction coefficient *α_ix,jy_* is weighted by the relative effort dedicated to foraging within its home location (1 *− f*). The second situation represents foraging of species *i* at location *x* on species *j* at location *y*. The net foraging effort *f* is equally divided among all locations reachable by species *i* from location *x* (the set given by *n_ix_*, which has a cardinality of *|n_ix_|*). The last situation is the opposite, where species *i* at location *x* is preyed upon by species *j* at location *y*. In this case, *f* is divided among all locations reachable by species *j* from location *y*. Note that this situation represents an equal division of foraging effort among all reachable locations.

Dispersal among different locations, in turn, is represented simply by the net variation in species densities between reachable locations, modelled by passive diffusion with coefficient *d* (Gravel et al., 2016):

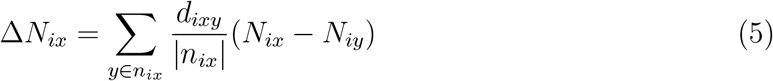

Thus, as it is the case with foraging, dispersal effort *d* is divided equally among all patches reachable by species *i* on location *x*.

## Parameterization and simulations

We considered predator-prey interactions, but the approach could easily be generalized to other types of interactions. The structure of local communities, i.e. who interacts with whom, is determined according to the niche model (Williams and Martinez, 2000), ensuring a realistig topology and that there are no disconnected species. We further assumed that the niche axis obtained from the niche model is linearly correlated with the foraging and dispersal distance of the different species, such that the species with lowest niche values could only forage or disperse to adjacent communities (Jacquet et al., 2017). Interaction coefficients at the regional scale are drawn from a normal distribution *N*(0.25, 0.1), with the sign structure given by the niche model. We introduced a small amount of spatial heterogeneity by drawing local coefficients from a normal distribution with mean centered on the corresponding regional coefficients and standard deviation of 0.1.

Local communities were placed along a single dimension space, which ends were connected together in order to maximize potential path lengths between non-connected communities, and prevent edge effects of communities at the end of the linear chain. We fixed the maximum dispersal and foraging distances to two cells away from the species’ home location in order to avoid excessive parameterization and for better comparing the net effects of the different movement types.

With this setting, we simulated the dynamics of 15 species at 10 local communities. The number of species and size of the landscape correspond to a meta-adjacency matrix of 15 * 10 = 150 rows. This size was chosen in order for the numerical calculation of the Jacobian matrices to be computationally feasible. Although a relatively small species richness and number of patches, it is sufficient to explore spatial distances and patch lengths of over 5 units.

We generated three sets of simulations: only dispersal, in which we set the dispersal coefficient *d* = 0.5, and the foraging coefficient *f* = 0 for every species; only foraging, with *d* = 0 and *f* = 0.5; and dispersal and foraging, with *d* = 0.5 and *f* = 0.5.

We ran 100 random topologies from each configuration, and for each replicate we obtained numerically the direct and net interaction coefficients between each pair of populations in the metacommunity, which could then be (1) qualitatively compared and (2) analyzed with regards to the distance between the interacting populations. We also computed basic quantitative descriptors of the direct and net effects networks at equlibrium: *connectance*, *average path length*, i.e. the average of the shortest path lengths between any pair of populations in the metacommunity, and *modularity*, which measures the tendency for nodes to be grouped into distinct modules (Newman, 2006). We calculated a weighted version of modularity that considers both positive and negative link weights, as implemented in the R *igraph* package (Csardi and Nepusz, 2006).

## Results

### What is the distribution of signs and magnitudes of net effects in communities connected by dispersal, foraging, or a mixture of both?

Net effects are mainly of equal sign to direct effects when local communities are connected by dispersal (Table 1), whereas sign switches occur in around 50% of pairwise interactions with foraging or a mixture of movement types. The ratio of positive to negative net effects is maintained at values close to 1, meaning a similar number of positive and negative net effects for all configurations.

**Table 1:** Summary metrics of the simulations performed. For each simulation, we group the results by location, i.e. whether the interaction occurs between species of the same (intra-) or different (inter-) patch. We show the ratio of positive to negative net effects, and the relative frequency of pairwise interactions that switch sign from direct to net effect.

| movement mode | location | +/- ratio | sign switches |  |
| --- | --- | --- | --- | --- |
|  |  |  | + to - | - to + |
| dispersal | intra-patch | 0.95 | 0.07 | 0.33 |
|  | inter-patch | 1.03 | 0.002 | 0 |
| foraging | intra-patch | 1.02 | 0.51 | 0.47 |
|  | inter-patch | 0.97 | 0.42 | 0.41 |
| dispersal and foraging | intra-patch | 1.02 | 0.52 | 0.57 |
|  | inter-patch | 1.02 | 0.48 | 0.49 |

The two types of movement and their combination displayed distinctly different net effects on interactions occurring both within the same location (intra-patch) and across different locations (inter-patch) (Fig. 2). Intraspecific effects across locations are generally positive with dispersal (blue points on the upper panel of Fig. 2). Intraspecific effects are however more variable and have a higher frequency of negative magnitudes with foraging and mixed movements (Fig. 2 middle and lower panels). Interspecific net effects are also generally of the same sign in local patches and across patches when communities are connected by dispersal (orange points on the upper panel of Fig. 2). Again, this trend is diluted with foraging in which case intra-patch and inter-patch interspecific effects display any combination of positive and negative signs, with no clear trend. Although here we analyze the results for *d* = 0.5 and *f* = 0.5, the distinctiveness of the effects of dispersal and foraging is maintained across a range of parameters (Appendix S1).

**Figure 2:**
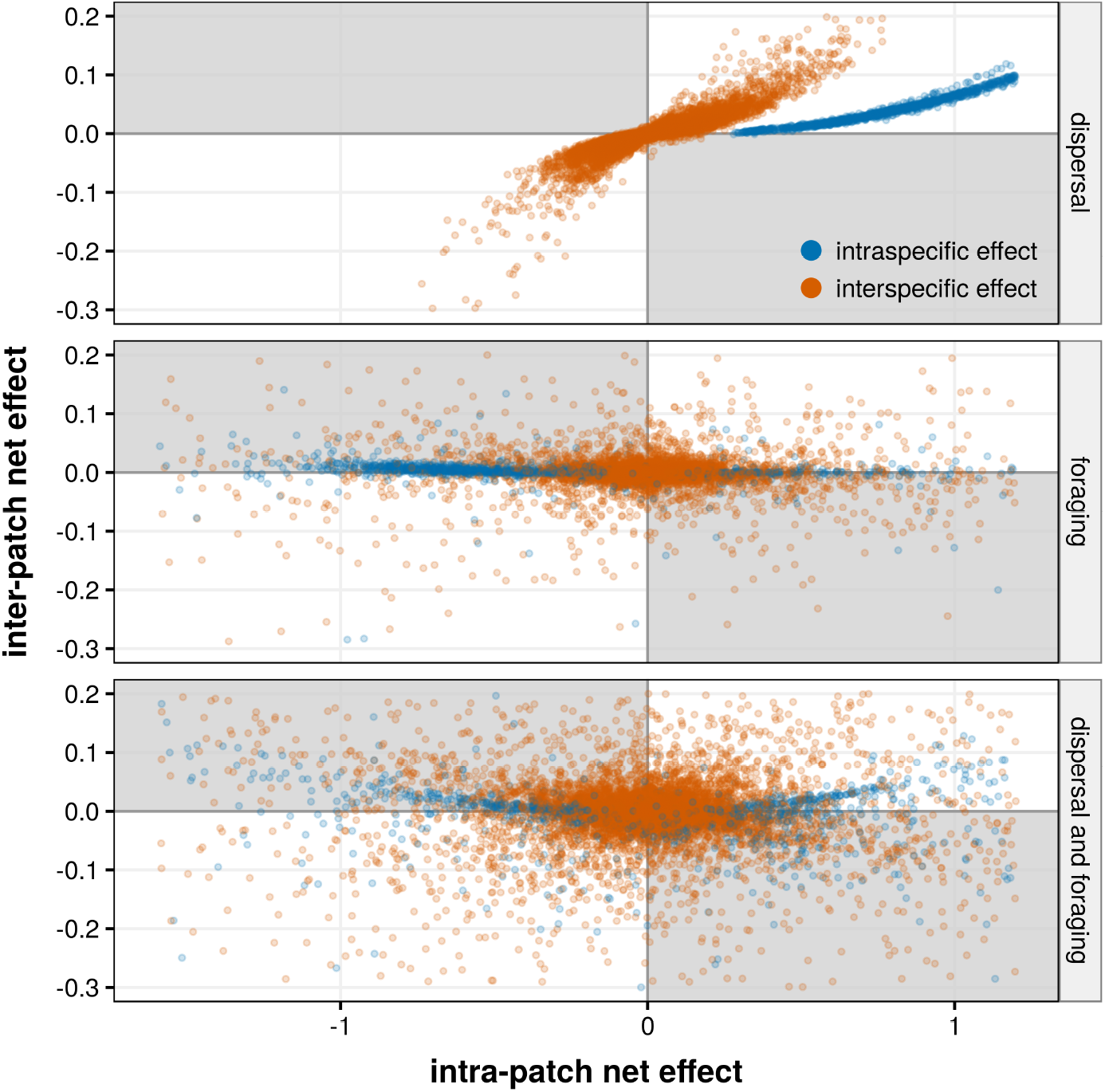
Distribution of intra and inter-patch net effects in the three configurations. Shaded quadrants are those where a sign switch occur between intra and inter-patch effects.

### Are networks of net effects similar in structure to networks of direct effects?

Virtually all pairs of species interact indirectly, as evidenced by net effect networks having connectances and path lengths of 1 in all cases (Fig. 3). In contrast, direct effect networks are obviously not fully connected, displaying intra-patch connectances at steady state of 0.38 on average and inter-patch connectances between 0.01 (dispersal only) and 0.1 (dispersal and foraging). Weighted modularity is also much higher in the direct effects networks than in the net effects ones, as expected. However, the modularity of the net effects networks also shows a decreasing trend from dispersal only networks to dispersal and foraging ones (Fig. 3).

**Figure 3:**
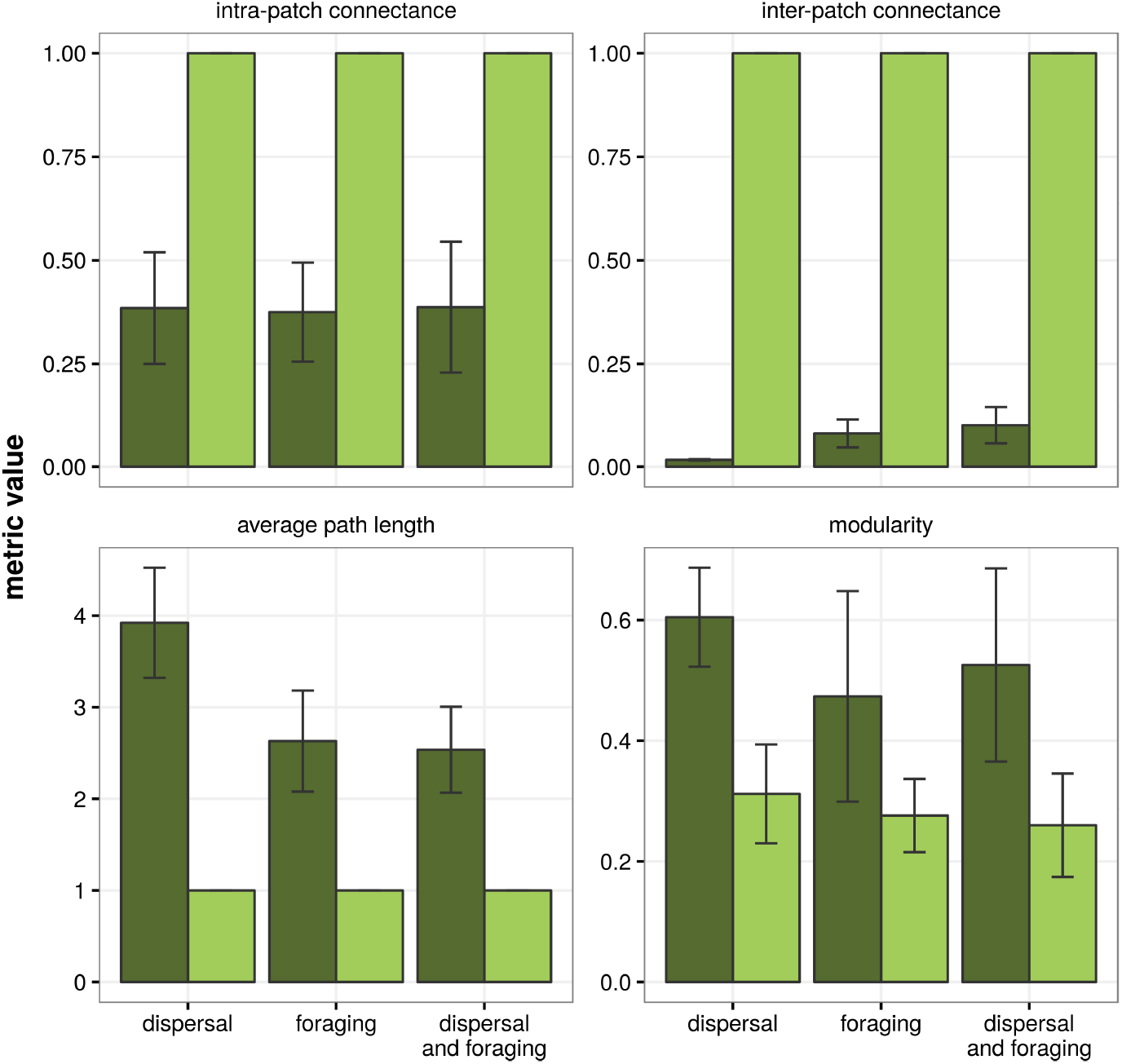
Network metrics of the metacommunities at equilibrium. Dark shades represent networks of direct effects, light shades networks of net effects, and error bars intervals of one standard deviation around the mean.

### Does the magnitude of the net effects between any two species decay with increasing distance between them?

The magnitude of spatial cascades is influenced by the length of its associated food chain. Net effects between any pair of species decrease in magnitude with the spatial distance between the two species (i.e. the distance between their home locations, assuming that connected locations are at distance 1 from each other). This decrease is even sharper when the distance metric considered is the path length between the two species, i.e. the number of connections separating them (Fig. 4). This result is observed with the two movement types and their combination, although the decay rate is generally highest with foraging. The trend is also robust to variations on dispersal and foraging rates (Appendix S1).

**Figure 4:**
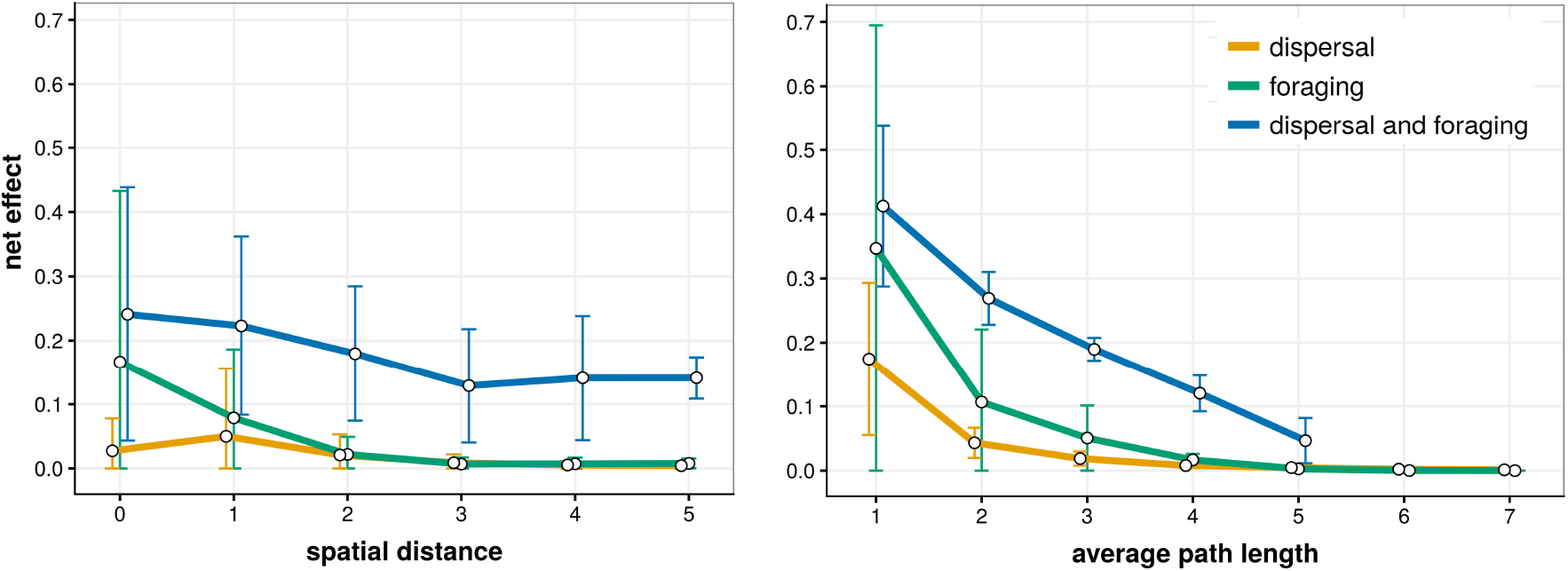
Net effect between species pairs as a function of *a)* spatial distance between patches, *b)* average path length between species. Values are averaged over all replicates, and error bars represent intervals of one standard deviation around the mean.

## Discussion

The importance of indirect effects, including trophic cascades, in driving ecosystem dynamics and structure is well established in theory (Abrams, 1992; Wootton, 2002; Gravel et al., 2010; Mayfield and Stouffer, 2017) and on the field (Menge, 1995; Peacor and Werner, 1997; Moya-Laraño and Wise, 2007; Barbosa et al., 2017; Trussell et al., 2017). The propagation of these effects across space has however not been studied systematically, despite many scattered observations of spatially explicit trophic cascades (Polis et al., 1997; Estes et al., 1998; Knight et al., 2005; Casini et al., 2012; Springer et al., 2018). Here we lay down basic tenets on how interaction effects are propagated in space depending on whether species connect different local communities via dispersal, foraging, or both modes of movement. We show that model metacommunities with populations connected by dispersal and foraging differ on (1) the proportion of pairwise interactions that switch sign between their direct and net effects, and (2) the sign and magnitudes of net effects on the local patch and across patches. Furthermore, the networks of net effects are markedly different from the direct effect ones, in all cases. Lastly, we observed that, in most cases, the magnitude of net effects between any two populations decays significantly with the distance between the two populations. In particular, the strongest decay occurs when distance is measured as the number of spatial connections necessary for linking the two populations. Indirect effects may generate unexpected net interaction outcomes between pairs of species. For example, Montoya et al. (2009) analyzed a set of well-resolved empirical food webs and showed that the influence of indirect effects induced a switch in interaction signs from direct to net effect for approximately 40% of species pairs. Using a similar approach, we show that net effects between populations of spatially disconnected communities may also be primarily driven by indirect feedbacks, as for example, cases in which a species foraging on a secondary location indirectly benefits its prey on this second location by altering the dynamics of the whole food web. This theoretical possibility has, to our knowledge, not been tested in empirical systems.

The “signatures” of net effects produced by dispersal, foraging, or mixed types of movement across localities are clearly different from each other (Amarasekare, 2008) even though we imposed equal maximum dispersal and foraging distances, and after accounting for different values of dispersal and foraging rates (Appendix S1). In the dispersal scenario, the almost complete concordance between the sign of intra- and inter-patch net effects points to a relatively homogeneous role of species at local and regional scales. Therefore, in natural systems with the same species pool connected mainly by dispersal, a local evaluation of the influence of the dispersing species may offer insights for the whole metacommunity. In the foraging or mixed modes, however, interspecific net effects derived from the populations of the same and different locations often switch signs, without showing any clear trend (middle and lower panels of Fig. 2). Therefore, within the assumptions of our model, when foraging is a prevalent mode of movement across locations, interspecific effects between any two species cannot be extrapolated from the local to the regional scale. In other words, a consumer that decreases a prey locally may have a positive effect on the same prey at the regional scale. Note, however, that our baseline scenario represents communities within an homogeneous habitat, the same species pool in all locations, with no density-dependent movement modes, no active selection or other types of heterogeneity. Furthermore, we deliberately chose dispersal and foraging modes of movement with identical maximum distances and temporal dynamics, in order to highlight their intrinsic differences. In reality, of course, both dispersal and foraging have extremely variable spatiotemporal scales. Foraging, in general, happens much faster than local demographic dynamics, which has led to characterize its effects as *spatial coupling* of local communities (Massol et al., 2011). This spatial coupling is thought to dampen population oscillations at lower trophic levels (McCann et al., 2005). The effects of dispersal, on the other hand, occur on temporal scales comparable to those of local dynamics, favouring different types of coexistence relationships, such as source-sink dynamics. The relative scales of foraging and dispersal are very heterogeneous, so that the spatial signal of interaction cascades will likely be correlated, in general, with these movement-related traits. In our model, we restricted the maximum dispersal and foraging rates to two cells away from the local community, and different values will presumably alter the net effect decay with spatial distance (Fig. 3 left panel). Interestingly, the decay of net effects with path length (Fig. 3, right panel) did not vary strongly with increases in maximum movement rates in our model (*d* and *f*, see Appendix S1). This result suggests that indirect effects may dampen the propagation of strong interactions across space, at least in environments with similar habitats and species pools.

Networks of net effects are fully connected (Fig. 3), meaning that every species has a non-zero influence in every other species of the metacommunity, through direct and/or indirect pathways, although the magnitude of this effect decays consistently with the distance and number of connections between species (Fig. 4). At the scale of local communities, Williams et al. (2002) reported that the majority of species in empirical food webs are separated by no more than two trophic links, and we have shown that in small metacommunities, this distance increases only slightly, to no more than four links on average (average path lengths of Fig. 3). The number of connections between local communities increases from dispersal to foraging modes, and is maximal when both modes are combined. This is reflected in the weighted modularity of the net effects networks, which tends to decrease along that axis. In the simple metacommunities modelled here, these structural patterns of the net effect networks are mostly confirmatory, but such analyses, when feasible, should provide novel insights into the structure and dynamics of more complex communities and metacommunities.

Dispersal generally represents unidirectional organismal movement across localities, with no associated direct movement of material. Foraging, on the other hand, involves a processing of organic material from the foraged location to the home location (Gounand et al., 2018). This two-way coupling has consequences for the role of the different locations as demographic sources or sinks (Gravel et al., 2010). In general, although here we have focused on movements of organisms across localities, the physical transfer of material often plays a key role in the structure and dynamics of the local communities (Polis et al., 1997). For example, the nutrients accumulated by Pacific salmon in open waters are transported to freshwaters during the spawning season, indirectly affecting terrestrial plants and animal populations (Naiman et al., 2002; Levi et al., 2012). The integration of these *ecological subsidies* with organismal movement is well develeoped theoretically in the framework of metaecosystem theory (Loreau et al., 2003; Gounand et al., 2018). We have provided a first approach to the explicit modelling of foraging behaviours in metacommunities, but its coupling with material transfers will expand and refine the results presented here. Most empirical studies on ecological subsidies are focused on transfers between different habitats, which usually involve specialized interactions (e.g. the consumption of Pacific salmon by grizzly bears) or species with life stages in different habitats, such as arthropods with aquatic larval stages that, in their adult form, switch to a predator role in terrestrial habitats. We, in turn, modelled communities with a small amount of spatial heterogeneity but the same pool of species and interactions on each patch. It is likely that interactions connecting different habitats are more specialized and have greater indirect effects than the general foraging interactions modelled here (in what has been called *keystone interactions*, Helfield and Naiman 2006). This specificity is likely to alter both the structure of the overall meta-network (Fig. 3) and the distance decay curves observed in our model system (Fig. 4). For example, Knight et al. (2005) showed how predatory fish could have strong net effects on terrestrial plants through a series of specialized interactions, even though the number of links separating these species is four (fish-dragonfly larvae-dragonfly adults-insect pollinators-plants).

Spatial distributions of net effects can also vary from our baseline expectations depending on the distances covered by the foraging or dispersing species. Intercontinental migrations are an extreme example of this, where strong coupled effects occur between species separated by thousands of kilometers (Alerstam and Bäckman, 2018). Further, such migratory movements connect localities at different moments in time, rather than continuously. The effect of such temporal decoupling may provoke strong oscillatory dynamics between systems (Springer et al., 2018), although in general, the stability dynamics associated with migrations have received little attention to date. Overall, the interplay between the rates and distances of dispersal and foraging, and their relationship to the spatial decay of net effects, clearly needs more attention in theoretical models and empirical studies. For example, experimental mesocosms allowing spatial movement of certain species among them may be used to test the differential influence of foraging and dispersal on local dynamics.

## Conclusions

We have provided a theoretical basis to the study of spatial propagation of indirect effects across ecosystems. We have shown that the net effect patterns generated by dispersal and foraging movements are clearly different. Furthermore, the structure of the metacommunity networks is markedly different depending on whether one considers direct interactions or net effects between species. The decay of net effect magnitude with distance, in our model, is the only result common to all simulations performed. These results may shed light on the spread of interaction effects in patches of the same habitat type, such as forest patches inserted in agricultural or urban areas. Furthermore, they represent a baseline case for developing more complex scenarios, such as the effects of interaction spread (1) across different habitat types and species pools, or (2) when material fluxes are accounted for.

## Acknowledgements

D.G.C. was funded by the Spanish Ministry of Education (FPU fellowship reference 2013/02147). M.B.A. acknowledges support from AAG-MAA/3764/2014 and CGL2015-68438-Pprojects

## References

Abrams, P. A. 1987. On Classifying Interactions between Populations. Oecologia 73:272–281.

Abrams, P. A. 1992. Predators That Benefit Prey and Prey That Harm Predators: Unusual Effects of Interacting Foraging Adaptation. The American Naturalist 140:573–600.

Alerstam, T., and J. Bäckman. 2018. Ecology of Animal Migration. Current Biology 28:R968–R972.

Allen, D. C., and J. S. Wesner. 2016. Comparing Effects of Resource and Consumer Fluxes into Recipient Food Webs Using Meta-Analysis. Ecology 97:594–604.

Amarasekare, P. 2008. Spatial Dynamics of Foodwebs. Annual Review of Ecology, Evolution, and Systematics 39:479–500.

Balcan, D., V. Colizza, B. Gonçalves, H. Hu, J. J. Ramasco, and A. Vespignani. 2009. Multiscale Mobility Networks and the Spatial Spreading of Infectious Diseases. Proceedings of the National Academy of Sciences 106:21484–21489.

Barbosa, M., G. W. Fernandes, O. T. Lewis, and R. J. Morris. 2017. Experimentally Reducing Species Abundance Indirectly Affects Food Web Structure and Robustness. Journal of Animal Ecology 86:327–336.

Barthélemy, M. 2011. Spatial Networks. Physics Reports 499:1–101.

Bender, E. A., T. J. Case, and M. E. Gilpin. 1984. Perturbation Experiments in Community Ecology: Theory and Practice. Ecology 65:1–13.

Berlow, E. L., A.-M. Neutel, J. E. Cohen, P. C. de Ruiter, B. Ebenman, M. C. Emmerson, J. W. Fox, V. A. A. Jansen, J. Iwan Jones, G. D. Kokkoris, D. O. Logofet, A. J. McKane, J. M. Montoya, and O. L. Petchey. 2004. Interaction Strengths in Food Webs: Issues and Opportunities. Journal of Animal Ecology 73:585–598.

Casini, M., T. Blenckner, C. Möllmann, A. Gårdmark, M. Lindegren, M. Llope, G. Kornilovs, M. Plikshs, and N. C. Stenseth. 2012. Predator Transitory Spillover Induces Trophic Cascades in Ecological Sinks. Proceedings of the National Academy of Sciences 109:8185–8189.

Chalfoun, A. D., F. R. Thompson, and M. J. Ratnaswamy. 2002. Nest Predators and Fragmentation: A Review and Meta-Analysis. Conservation Biology 16:306–318.

Christian, C. E. 2001. Consequences of a Biological Invasion Reveal the Importance of Mutualism for Plant Communities. Nature 413:635–639.

Csardi, G., and T. Nepusz. 2006. The Igraph Software Package for Complex Network Research. InterJournal Complex Systems:1695.

Estes, J. A., M. T. Tinker, T. M. Williams, and D. F. Doak. 1998. Killer Whale Predation on Sea Otters Linking Oceanic and Nearshore Ecosystems. Science 282:473–476.

Gounand, I., E. Harvey, C. J. Little, and F. Altermatt. 2018. Meta-Ecosystems 2.0: Rooting the Theory into the Field. Trends in Ecology & Evolution 33:36–46.

Gravel, D., F. Guichard, M. Loreau, and N. Mouquet. 2010. Source and Sink Dynamics in Meta-Ecosystems. Ecology 91:2172–2184.

Gravel, D., F. Massol, and M. A. Leibold. 2016. Stability and Complexity in Model Meta-Ecosystems. Nature communications 7:12457.

Helfield, J. M., and R. J. Naiman. 2006. Keystone Interactions: Salmon and Bear in Riparian Forests of Alaska. Ecosystems 9:167–180.

Jacquet, C., D. Mouillot, M. Kulbicki, and D. Gravel. 2017. Extensions of Island Biogeography Theory Predict the Scaling of Functional Trait Composition with Habitat Area and Isolation. Ecology Letters 20:135–146.

Knight, T. M., M. W. McCoy, J. M. Chase, K. A. McCoy, and R. D. Holt. 2005. Trophic Cascades across Ecosystems. Nature 437:nature03962.

Leibold, M. A., and J. M. Chase. 2018. Metacommunity Ecology. Number 59 in Monographs in population biology, Princeton University Press, Princeton.

Leibold, M. A., M. Holyoak, N. Mouquet, P. Amarasekare, J. M. Chase, M. F. Hoopes, R. D. Holt, J. B. Shurin, R. Law, D. Tilman, M. Loreau, and A. Gonzalez. 2004. The Metacommunity Concept: A Framework for Multi-Scale Community Ecology. Ecology Letters 7:601–613.

Levi, T., C. T. Darimont, M. MacDuffee, M. Mangel, P. Paquet, and C. C. Wilmers. 2012. Using Grizzly Bears to Assess Harvest-Ecosystem Tradeoffs in Salmon Fisheries. PLOS Biology 10:e1001303.

Loreau, M., N. Mouquet, and R. D. Holt. 2003. Meta-Ecosystems: A Theoretical Framework for a Spatial Ecosystem Ecology. Ecology Letters 6:673–679.

Massol, F., D. Gravel, N. Mouquet, M. W. Cadotte, T. Fukami, and M. A. Leibold. 2011. Linking Community and Ecosystem Dynamics through Spatial Ecology. Ecology Letters 14:313–323.

Mayfield, M. M., and D. B. Stouffer. 2017. Higher-Order Interactions Capture Unexplained Complexity in Diverse Communities. Nature Ecology & Evolution 1:0062.

McCann, K. S., J. B. Rasmussen, and J. Umbanhowar. 2005. The Dynamics of Spatially Coupled Food Webs. Ecology Letters 8:513–523.

Menge, B. A. 1995. Indirect Effects in Marine Rocky Intertidal Interaction Webs: Patterns and Importance. Ecological Monographs 65:21–74.

Montoya, J. M., G. Woodward, M. C. Emmerson, and R. V. Solé. 2009. Press Perturbations and Indirect Effects in Real Food Webs. Ecology 90:2426–2433.

Moya-Laraño, J., and D. H. Wise. 2007. Direct and Indirect Effects of Ants on a Forest-Floor Food Web. Ecology 88:1454–1465.

Naiman, R. J., R. E. Bilby, D. E. Schindler, and J. M. Helfield. 2002. Pacific Salmon, Nutrients, and the Dynamics of Freshwater and Riparian Ecosystems. Ecosystems 5:399–417.

Newman, M. E. J. 2006. Modularity and Community Structure in Networks. Proceedings of the National Academy of Sciences 103:8577–8582.

Novak, M., J. D. Yeakel, A. E. Noble, D. F. Doak, M. Emmerson, J. A. Estes, U. Jacob, M. T. Tinker, and J. T. Wootton. 2016. Characterizing Species Interactions to Understand Press Perturbations: What Is the Community Matrix? Annual Review of Ecology, Evolution, and Systematics 47:409–432.

Orians, G. H. 1979. On the Theory of Central Place Foraging. Analysis of ecological systems pages 157–177.

Orrock, J. L., J. H. Grabowski, J. H. Pantel, S. D. Peacor, B. L. Peckarsky, A. Sih, and E. E. Werner. 2008. Consumptive and Nonconsumptive Effects of Predators on Metacommunities of Competing Prey. Ecology 89:2426–2435.

Peacor, S. D., and E. E. Werner. 1997. Trait-Mediated Indirect Interactions in a Simple Aquatic Food Web. Ecology 78:1146–1156.

Polis, G. A. 1994. Food Webs, Trophic Cascades and Community Structure. Australian Journal of Ecology 19:121–136.

Polis, G. A., W. B. Anderson, and R. D. Holt. 1997. Toward an Integration of Landscape and Food Web Ecology: The Dynamics of Spatially Subsidized Food Webs. Annual Review of Ecology and Systematics 28:289–316.

Schmitz, O. J., P. A. Hambäck, and A. P. Beckerman. 2000. Trophic Cascades in Terrestrial Systems: A Review of the Effects of Carnivore Removals on Plants. The American Naturalist 155:141–153.

Springer, A. M., G. B. van Vliet, N. Bool, M. Crowley, P. Fullagar, M.-A. Lea, R. Monash, C. Price, C. Vertigan, and E. J. Woehler. 2018. Transhemispheric Ecosystem Disservices of Pink Salmon in a Pacific Ocean Macrosystem. Proceedings of the National Academy of Sciences 115:E5038–E5045.

Swihart, R. K., N. A. Slade, and B. J. Bergstrom. 1988. Relating Body Size to the Rate of Home Range Use in Mammals. Ecology 69:393–399.

Trussell, G. C., C. M. Matassa, and P. J. Ewanchuk. 2017. Moving beyond Linear Food Chains: Trait-Mediated Indirect Interactions in a Rocky Intertidal Food Web. Proceedings of the Royal Society B: Biological Sciences 284:20162590.

Williams, R. J., E. L. Berlow, J. A. Dunne, A.-L. Barabási, and N. D. Martinez. 2002. Two Degrees of Separation in Complex Food Webs. Proceedings of the National Academy of Sciences 99:12913–12916.

Williams, R. J., and N. D. Martinez. 2000. Simple Rules Yield Complex Food Webs. Nature 404:180–3.

Wootton, J. T. 2002. Indirect Effects in Complex Ecosystems: Recent Progress and Future Challenges. Journal of Sea Research 48:157–172.

Zhao, L., H. Zhang, E. J. O’Gorman, W. Tian, A. Ma, J. C. Moore, S. R. Borrett, and G. Woodward. 2016. Weighting and Indirect Effects Identify Keystone Species in Food Webs. Ecology Letters 19:1032–1040.

